# A human-aware control paradigm for human-robot interactions, a simulation study

**DOI:** 10.1101/2024.03.20.585749

**Authors:** Reza Sharif Razavian

## Abstract

This paper presents a novel model for predicting human movements and introduces a new control method for human-robot interaction based on this model. The developed predictive model of human movement is a *holistic* model that is based on well-supported neuroscientific and biomechanical theories of human motor control; it includes multiple levels of the human sensorimotor system hierarchy, including high-level decision-making based on internal models, muscle synergies, and physiological muscle mechanics. Therefore, this holistic model can predict arm kinematics and neuromuscular activities in a computationally efficient way. The computational efficiency of the model also makes it suitable for repetitive predictive simulations within a robot’s control algorithm to predict the user’s behavior in human-robot interactions. Therefore, based on this model and the nonlinear model predictive control framework, a *human-aware control* algorithm is implemented, which internally runs simulations to predict the user’s interactive movement patterns in the future. Consequently, it can optimize the robot’s motor torques to minimize an index, such as the user’s neuromuscular effort. Simulation results of the holistic model and its utilization in the human-aware control of a two-link robot arm are presented. The holistic model is shown to replicate salient features of human movements. The human-aware controller’s ability to predict and minimize the user’s neuromuscular effort in a collaborative task is also demonstrated in simulation.

## 1 INTRODUCTION

As robots become more integrated into human environments, ensuring that they can interact with people safely and intuitively becomes increasingly important. This is a challenging control problem that is especially critical in physical human-robot interactions.

In control systems, high-quality performance is only possible if the dynamics of the system being controlled is adequately modeled. This also applies to the control of human-robot interactions—To control the coupled human-robot dynamics, it is essential to model the entire interaction, including the user’s movements. For example, in wearable robots, exoskeletons, surgical robots, and industrial collaborative robots (cobots), where the human user and the robot share a physical task, the robot must be able to predict the user’s behavior and act in a predictive rather than reactive way. To achieve this predictive control, mathematical models must be developed to predict human movements based on minimal information, such as the task’s high-level goals. Additionally, these models must be able to run many times within a single cycle of the robot’s controller to allow the robot to compare predictions and choose the best course of action.

Researchers in the fields of biomechanics and motor neuroscience have sought to present such predictive models for human movements, and significant achievements have been made. In musculoskeletal biomechanics, and especially gait biomechanics, large-scale nonlinear musculoskeletal models that rely on nonlinear optimization algorithms have demonstrated great success in predicting movement kinematics [1], muscle activity patterns [2], joint contact forces [3], and human-machine interaction [4] (see the review papers [5–7] for more success stories.) These biomechanical models are rich in details, such as musculoskeletal anatomy and mechanical properties of the limbs; however, the downside of these models is that they are slow to run—it may take minutes to hours to predict a few seconds of movement. Therefore, the computational efficiency of these models is a limiting factor that prevents them from being used for real-time predictive control applications.

On the other end of the model complexity spectrum, the neuroscientific models of movement control seek to trim as much detail as possible and present a minimalistic model that captures salient features of human motor control. A prominent example of such models is based on stochastic linear optimal feedback control theory [8, 9], which provides a mechanistic explanation of how an *internal model* of the world is used to process sensory information and plan and execute movements. Despite their simplicity, these neuroscientific models are invaluable since they explain *why* we move the way we do. Therefore, these models succeed at an objective that is different from that of biomechanical models. The former discusses the “why”, while the latter focuses on the “how.”

The challenge is that human movements are the emerging behavior of a complex dynamical system that constitutes the neural, muscular, skeletal, and sensory systems, which interact with the environment in a closed-loop manner. To have a truly predictive model, we need to take into account both the high-level (the why) and low-level (how) details in a *holistic* model.

In this paper, I present a novel model for human movement with these considerations in mind. This *holistic* model encompasses multiple levels of detail in the human sensorimotor system; it is based on neuroscientific theories such as internal models and optimal sensorimotor integration for decision-making and action planning, as well as biomechanically plausible details of the musculoskeletal system. Therefore, this model enables predicting and distinguishing the contributions of various levels of the sensorimotor system to the user’s behavior.

In addition to the model, I also present the preliminary results of a new control paradigm for human-robot interaction, called *human-aware control*. This controller, which is based on nonlinear model predictive control, employs the developed holistic model of human movement to predict the user’s behavior during the interactions and optimize the robot’s joint torques in real-time. This control paradigm is advantageous over the existing impedance/admittance-based controllers, e.g., [10–12], since the robot no longer treats the interaction as a disturbance to be observed and compensated.

## 2 Methods

The human-aware control paradigm (Figure 1) consists of two major components: (1) a holistic model for human movement, which allows the robot controller to predict how the user reacts to the robot’s actions, and (2) an optimal controller, based on nonlinear model predictive control, which employs the holistic model to predict the future behavior of the user and optimize motor torques.

**Figure 1.**
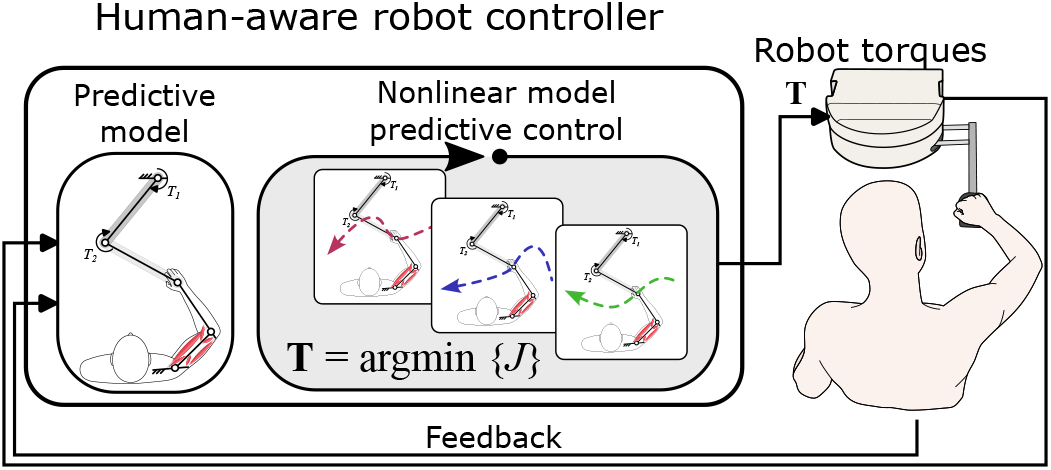
The human-aware control of robot based on the nonlinear model predictive control scheme. A holistic model of human movement predicts the user’s behavior. The model is run iteratively to find the best robot actions.

### 2.1 Holistic model of human movements

Despite the nonlinearities and redundancies present in the musculoskeletal system, the nervous system manages to control human movements dexterously and in real time. Therefore, replicating the same control principles *in silico* is the best strategy to predict human movements in a computationally efficient and biologically plausible way. To this end, a holistic model for human movements is developed (Figure 2), which is based on existing theories in motor neuroscience and biomechanics. This holistic model has a hierarchical structure representing different “levels” of control in the nervous system, which work together cohesively in a close-loop manner (Figure 2**B**). The highest level of control (described in section 2.1.1) is an abstract computational module that deals with action planning and decision-making. These abstract decisions are made based on a low-dimensional *internal model* [13] that resides in a simplified and abstract task space. This module receives sensory feedback from the body and optimally infers the current states of its internal model. Using these estimated internal states, an optimally tuned feedback control gain calculates some abstracted muscle commands in the low-dimensional task space. This low-dimensional output is then expanded into high-dimensional physiological muscle space in a mid-level module (section 2.1.2) by utilizing coordinated muscle activity patterns, often known as *muscle synergies* [14]. Finally, the low-level biomechanical model (section 2.1.3), which represents the nonlinear dynamics of the muscles, arm, and the environment, receives muscle activities and produces movements in the physical world. The sensory information about the kinematics of the hand (the physical task space) is fed back to the higher-level modules to close the loop.

**Figure 2.**
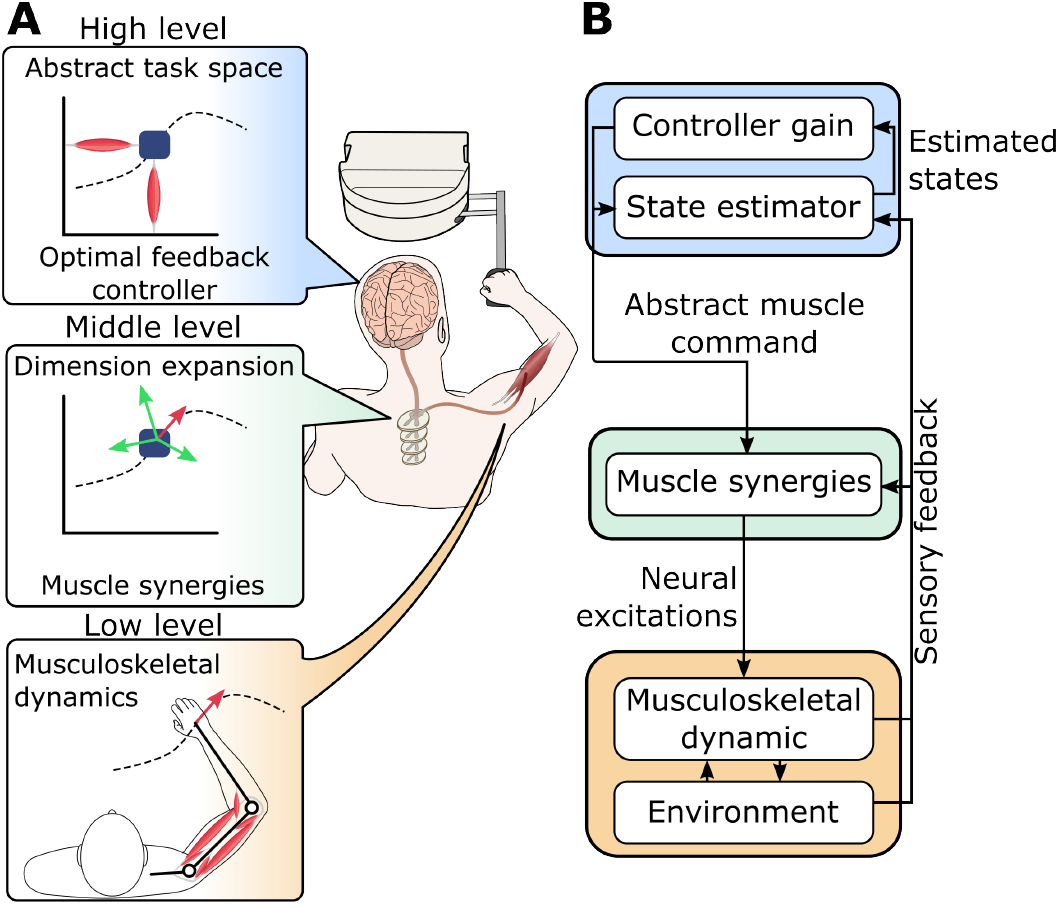
**A**. The overview of the holistic model of the human movement. Each module represents a distinct level in the sensorimotor control hierarchy. **B**. The hierarchical structure of the model based on biomechanical and neuroscientific theories and the information flow between modules.

#### 2.1.1 The high-level module: decision making based on internal models

This module is based on the mounting evidence in motor neuroscience research, which supports the argument that humans chose a movement strategy that minimizes a combination of effort and task accuracy [8, 9, 15]. These models often focus only on the “task-level” dynamics—they abstract the entire movement, such as reaching, into a point-mass being moved around in a 2- or 3-dimensional *task space*. Although these highly simplified models fall short of representing the entire complexity of the neuromuscular system, their success in replicating patterns of movement in the task space is a strong indicator of their validity as a fundamental mechanism for movement-related decision-making [16–18]. In addition to their validity, the computational simplicity of these models makes them an ideal choice for the high-level decision-making module in the holistic model.

This high-level module is an extension of linear quadratic Gaussian (LQG) control, which is adapted to the multiplicative characteristics of the noise in the sensorimotor system—the standard deviation of noise scales linearly with the amplitude of the signal [19, 20]. In this high-level controller, the *internal model*, i.e., the model based on which motor decisions are made, is a highly simplified 2-dimensional system consisting of a point-mass actuated by two abstract “muscles” (Figure 3). This abstract task space represents the 2-dimensional position of the hand in the physical space. The equations of motion for this internal model along the *x* and *y* dimensions are:

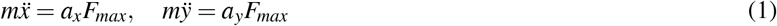

where *F*_*max*_ is the maximum muscle force that is scaled by the muscle activations *a*, which follow first-order dynamics:

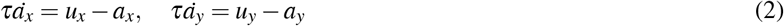

Here, *τ* is the muscles’ activation time constant, and *u*’s are the abstract neural excitation signals that the high-level controller determines. The numerical values of all parameters are provided in Table 1.

**Figure 3.**
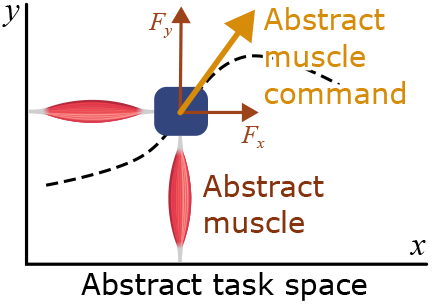
The *internal model* used in the high-level controller. The model is in the abstract 2-dimensional task space that represents hand position. Two abstract muscles move a point mass in this space; the total muscle force is an *abstract muscle command* that is sent to the lower-level module.

**Table 1:** Numerical values of the parameters in the simulations.

| Internal model |  |  |  |
| --- | --- | --- | --- |
| Parameter | Value | Parameter | Value |
| $m$ | 3 kg | $\xi$ | $\text{diag}_{6 \times 6}(2 \times 10^{-7})$ |
| $\tau$ | 30 ms | $\omega$ | $\text{diag}_{6 \times 6}(10^{-6})$ |
| $F_{max}$ | 1000 N | $\mathbf{C}$ | $\text{diag}_{2 \times 2}(2 \times 10^{-1})$ |
| $\mathbf{R}$ | $\text{diag}_{2 \times 2}^*(1)$ | $\eta$ | $\text{diag}_{2 \times 2}(10^{-9})$ |
| $\mathbf{Q}_{k < N-D}$ | $\text{diag}_{6 \times 6}(0)$ | $d$ | 50 ms |
| $\mathbf{Q}_{k \geq N-D}$ | $\text{diag}_{6 \times 6}(1)$ | | |
| Musculoskeletal model |  |  |  |
| See section 4.1.6 in [32] |  |  |  |
| Robot model |  |  |  |
| Parameter | Value | Description |  |
| $l_1 = l_2$ | 0.3 m | Length of each link | |
| $m_1 = m_2$ | 0.5 kg | Mass of each link | |
| $d_1 = d_2$ | 0.55 m | Distance to center of mass | |
| $I_1 = I_2$ | $5 \times 10^{-3} \text{ kg.m}^2$ | Moment of inertia about center of mass | |
| Human-aware robot controller |  |  |  |
| Parameter | Value | Description |  |
| $\Delta t$ | 5 ms | Discretization step size | |
| $N_p$ | 50 time steps | Prediction horizon length | |
| $D$ | 21 time steps | Dwell time at target | |
| $N$ | 221 time steps | Total duration of simulation | |

\* A diagonal matrix of given size with those parameters on the diagonal

To solve the high-level control problem, this internal model is transformed into a discrete state-space form

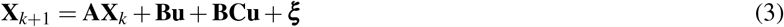

with 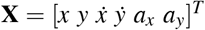 and **u** = [*u*_*x*_ *u*_*y*_]^*T*^ being the state and input vectors, respectively, and **A, B** being the corresponding discrete state and input matrices. The subscript *k* represents *k*^*th*^ time step. It is assumed that the dynamics of this internal model is subject to multiplicative noise (through **BCu**) and additive noise (***ξ***) with Gaussian distributions. These noise characteristics in the internal model are essential as they are shown to be important contributors to human behavior [19, 21].

The high-level control calculates the optimal inputs **u**^*∗*^ as

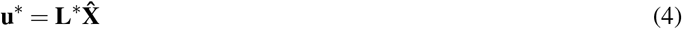

Here, **L**^*∗*^ is the optimally tuned feedback gain, and 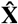 is the estimated state vector, which is inferred from the delayed and noisy sensory information **Y** using a Kalman filter

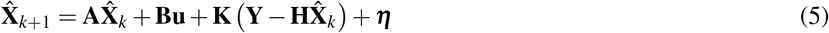

where **K** is the Kalman filter gain. This state estimation process is also assumed to be contaminated with additive Gaussian noise ***η***. To calculate the Kalman gain, the sensory information **Y** is assumed to be a full readout (**H** = **I**_6*×*6_) of the internal state vector that is contaminated with noise and delay:

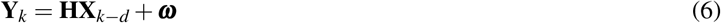

with *d* representing the sensory delay, and ***ω*** being Gaussian noise. However, during movements, the sensory information from the musculoskeletal model is used (see section 2.1.3).

Because of the multiplicative noise in the state-space equations (3), the optimal controller and Kalman gains must be obtained iteratively [20]. The control gain is found to minimize the quadratic cost function:

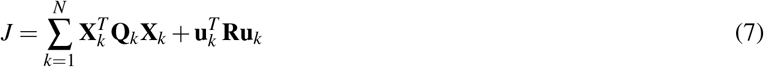

where **Q**_*k*_ and **R** are the state and control penalty weights. The control penalty **R** is constant, but the state penalty **Q**_*k*_ is time-dependent; it is zero throughout the motion and only contains non-zero elements along its main diagonal in the last *D* time steps (dwell time) to penalize the position and velocity at the target. In other words, the high-level controller seeks to reach the target within *N −D* time steps with minimal effort, but the trajectory is unspecified.

The estimated internal states in the abstract task space are considered to be the *perceived* motion; i.e., this is how the high-level controller (the “brain”) thinks the movement is progressing. Based on these perceived states of the world, the high-level controller calculates the abstract muscle commands to move the internal model toward the intended target. These low-dimensional muscle commands in the abstract task space must be translated into physiological muscle commands. This is done in the mid-level module.

#### 2.1.2 The mid-level module: Dimension expansion with muscle synergies

To be physiologically realistic, the high-level controller’s abstract and low-dimensional commands must be expanded into the high-dimensional muscle space. The first challenge is that physiological muscles can only pull, unlike the abstract muscles that can pull and push. Further, the human musculoskeletal arm has more muscles than degrees of freedom, leading to infinite possible solutions for muscle forces that can produce a specific task space force. Lastly, the nonlinear dynamics of the multi-link arm, as opposed to the linear dynamics of the point mass, must be taken into account to produce accurate movements. Therefore, solving for muscle forces that replicate the task-space movement as predicted by the internal model is not straightforward. This problem—referred to as the *muscle load-sharing problem* in biomechanics—is usually solved using nonlinear optimization [5, 6]; a process that is neither physiologically plausible nor real-time implementable. Instead, this holistic model employs a biologically plausible approach to solving muscle activities using the well-supported theory of *muscle synergies* [14].

The overview of the dimension expansion module is shown in Figure 4. In this approach, muscle synergies are defined as activating a group of muscles with known relative ratios [22] (Figure 4**A**). It is postulated that the nervous system holds a memory of synergies and their actions in the task space, i.e., the nervous system knows the hand force vectors produced by each synergy (Figure 4**B**). These synergy-produced force vectors are considered a *basis set* for the task space [23, 24]. The abstract muscle command in the low-dimensional task space is projected onto these basis vectors to calculate the coefficient of the force along each basis. Because usually there are more synergies (in this case 4) than the dimensions of the task space (*n* = 2), the decomposition solution is not unique; therefore, a non-negative least-squares algorithm is used to solve for the non-negative coefficient vector **C** in the equation

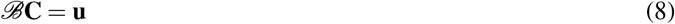

Here *ℬ*_2*×*4_ is the matrix formed by the 4 synergy-produced basis vectors in the 2-dimensional task space, and **u**_2*×*1_ is the abstract muscle command from the high-level controller. Once the coefficients *C*_*i*_ are calculated, the weighted sum of the synergies (Figure 4**C**) results in the “neural” excitation commands **e** that represent aggregate motoneuron activities.

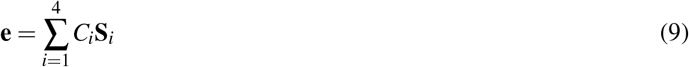

Here **S**_*i*_ is the vector of muscle activation weights in the *i*^*th*^ synergy.

**Figure 4.**
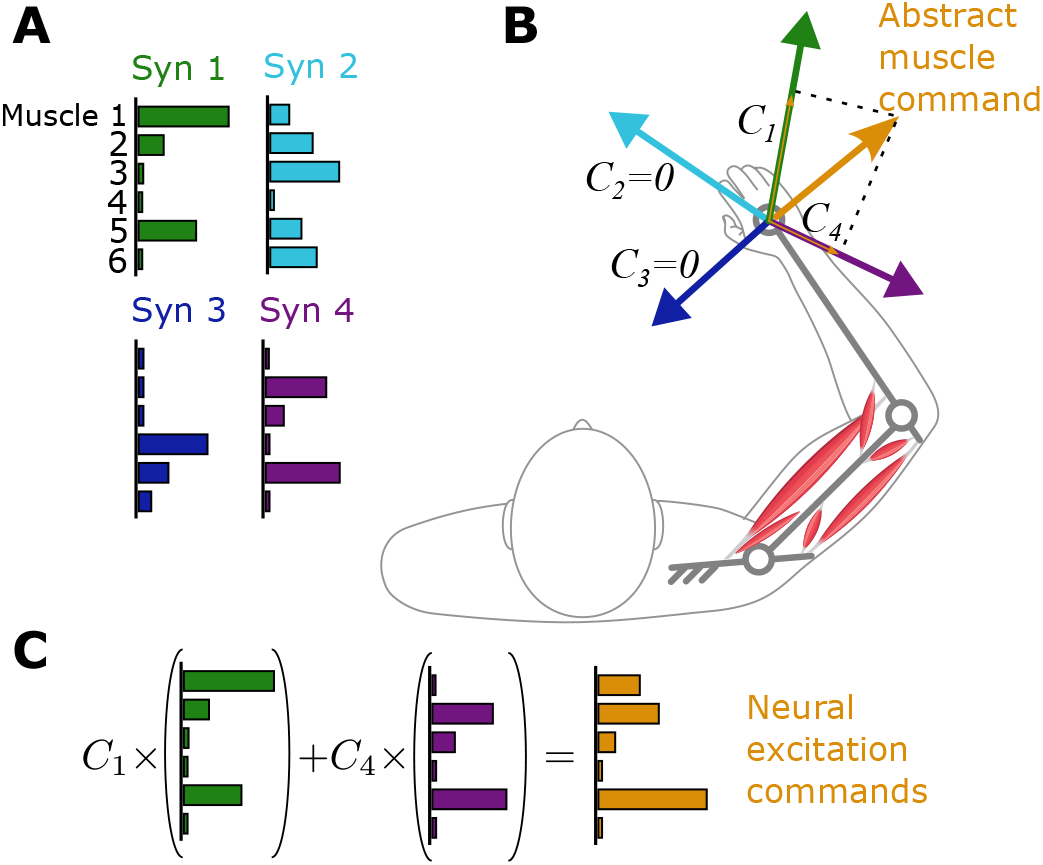
The mid-level dimension expansion module using muscle synergies. **A**. Synergies are co-activations of muscles with known relative ratios. **B**. The effect of each synergy in the task space is a force vector known to the motor controller. The low-dimensional neural excitation command in the task space is decomposed onto the synergy-produced bases to calculate the activation coefficient of each synergy, *C*’s. **C**. Synergies are combined with the calculated coefficients to produce neural excitation commands.

An offline optimization and data reduction procedure can be used to obtain the synergies. The details are available in [23,25] and are omitted here for brevity. This offline approach to learning and storing synergies parallels skill acquisition and motor memory formation in humans [26].

The calculated neural excitation commands can be directly used as the input to the musculoskeletal model (see section 2.1.3). However, because of the nonlinear dynamics of the multi-link arm, velocity-dependent accelerations lead to a task space motion that is different from the desired one. To overcome that, the high-level muscle commands are adjusted using the known dynamics of the arm to compensate for the velocity-dependent acceleration. For details, the readers are referred to [25].

The mid-level module’s output is a set of neural excitation commands *e* that are fed to the musculoskeletal model.

#### 2.1.3 The low-level module: Musculoskeletal and environment dynamics

In this study, the musculoskeletal model representing the body is a 2-link planar linkage that is actuated by 6 muscles (Figure 5). This model type is the standard upper extremity model for various biomechanical and motor control studies [27–30], as it captures the important features of a human musculoskeletal system: muscle redundancy, nonlinear muscle mechanics, and nonlinear dynamics of the arm. The two links in the model represent the upper arm and the forearm, which are connected to a fixed torso and to each other via revolute joints. The wrist is not modeled. The muscles are modeled as Hill-type muscles [31] that produce force *F* according to

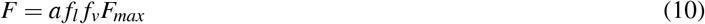

with *f*_*l*_ and *f*_*v*_ being the force-length and force-velocity relationship [31]. *a ∈* [0, 1] is the activation of the muscle, and *F*_*max*_ is maximum isometric muscle force. The muscle activation is driven by the neural excitation signal *e* according to the dynamics

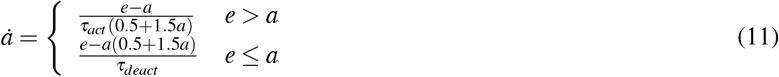

with *τ*_*act*_ and *τ*_*deact*_ being the muscles’ activation and deactivation time constants. The contractile forces (10) are applied to arm links along each muscle’s line of action, which is defined as the straight line connecting the origin and insertion points of the muscle. The tendons and other compliance of the muscle are neglected in this musculoskeletal model. The musculoskeletal model parameters are taken from [32].

**Figure 5.**
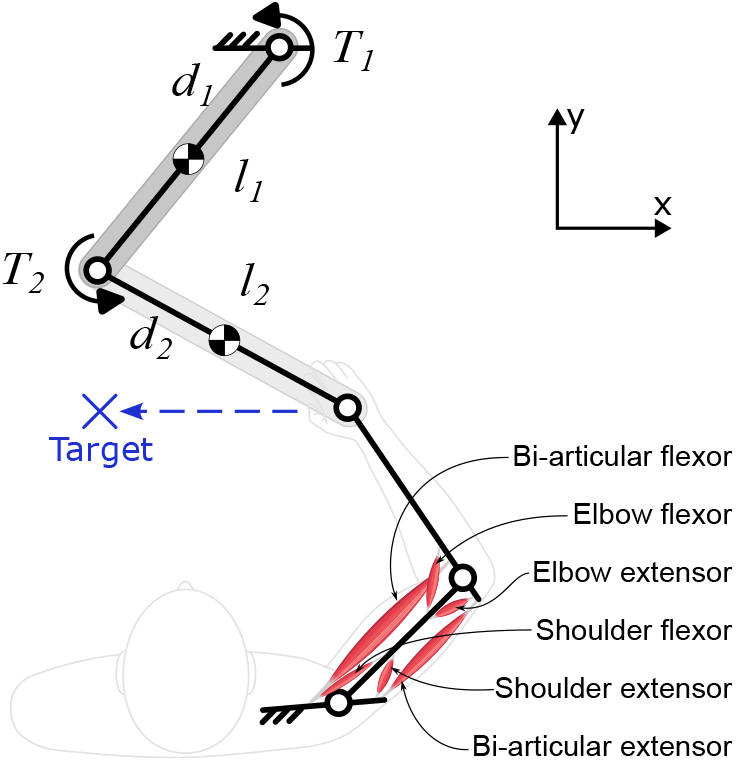
The musculoskeletal arm coupled with the 2-link robot

The sensory information from the musculoskeletal arm closes the control loop in the holistic model. The high-level controller receives the delayed feedback (with the same delay time *d* as in (6)) from the musculoskeletal hand position and velocity to estimate the internal model’s states according to (5). Therefore, the estimated internal states are updated based on the actual movement’s measurements rather than the internal model’s state dynamics.

In the human-robot interaction simulations, a revolute joint connects the “hand” of the musculoskeletal arm to the end-effector of a 2-link planar robotic arm (Figure 5). The robot’s controller (section 2.2) determines the joint torques *T* .

### 2.2 Human-aware control of robot

The human-aware control paradigm is based on a nonlinear model predictive control scheme [30, 33] (Figure 1). In this controller, the *control-oriented model* is the holistic model presented in the previous section. The robot’s controller can internally run this model to predict the user’s behavior for a short time into the future. It is assumed that the robot and its control-oriented model are aware of the goal of the task: to reach a known target in the task space (Figure 5). At a given time step during the motion, the human-aware controller initializes its control-oriented model using the current measurements from the plant and runs predictive simulations for the duration of the *prediction horizon N*_*p*_ into the future. The predictive simulation informs the controller how the human will continue the motion, given the interactive forces supplied by the robot. The controller calculates the two motor torques, *T*_1_ and *T*_2_, to minimize an objective computed over the prediction horizon. Then, the controller applies these optimal torques to the robot for one time step. The entire process repeats in the next time step with updated measurements from the system. The objective function may be a function of any variables in the control-oriented model, e.g., those in the high-, mid-, or low-level modules. The simple objective function considered in this study is the minimization of the user’s effort:

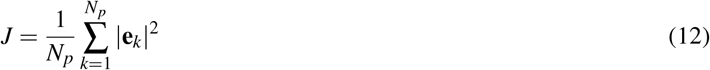

where **e**_*k*_ is the vector of neural excitations in the holistic model. The summation is over the time steps *k* from the beginning to the end of the prediction horizon with length *N*_*p*_. To have smooth torque profiles over the prediction horizon, the robot’s motor torques are parameterized using 2nd-order polynomial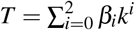, and the decision variables in this *receding horizon* optimization are the 6 coefficients *β*_*i*_ for the two robot torques. The controller uses a nonlinear optimization algorithm to find the robot’s joint torque coefficients such that the objective function (12) is minimized.

In the present implementation of the human-aware controller, it is assumed that the controller has access to the current measurement of the robot’s joint positions and velocities, as well as the human’s joint angle and velocities, muscle activities, and internal model’s states. It is not realistic to assume that the robot has a direct measurement of the human’s internal states, but in the future, nonlinear estate estimation techniques such as moving horizon estimator [34, 35] may be used to provide the controller with such information.

## 3 Simulation setting

The holistic model of movements and the human-aware control paradigm were implemented and evaluated in simulation. In the human-robot interaction simulation, the *plant* was the same holistic model discussed above. I.e., the plant model for the simulations was the same as the control-oriented model. The simulated task was to move the hand to a specified target 25 *cm* to the left within a prescribed time of 1 *s*. Four simulation settings were studied: *A. No interaction*: The human moved to the target without interacting with the robot. *B. Robot off* : The human moved in physical contact with the robot, but robot torques are set to zero. *C. Human-aware control*: The robot was controlled with the human-aware controller to assist with the task and minimize the user’s effort (12). *D. Impedance control*: The robot was controlled with an impedance controller [36] that was set to follow the reference trajectory obtained in the “no-interaction” scenario A. This impedance controller, being the standard control method for collaborative robots, was implemented as the baseline against which the optimal human-aware controller was compared.

The input to simulations A-C was only the *target position* and *movement duration*. The fourth simulation with the impedance controller also required the reference trajectory as the input, which was obtained from the first simulation. All simulations were done in MATLAB 2022b (Mathworks Inc., Natick, MA), with the numerical parameters as in Table 1.

## 4 Results

Four simulations were run to showcase and compare separate aspects of the holistic model and the human-aware controller.

The first simulation (Figure 6**A**) verified how the model could replicate human movement in the absence of interaction with the environment. The high-level movement decisions made based on the internal model (first two rows in Figure 6) could move the arm with kinematic trajectories that closely match unrestricted human movements with smooth transitions and bell-shaped velocity profiles [37]. Expanding the abstract motor commands, first into synergy activations and subsequently into physiological neural excitations (rows 3, 4), calculated muscle forces (row 5) that moved the musculoskeletal arm with the same kinematics in the physical task space (row 6). The sensory feedback sent to the high-level controller ensured accurate estimation of internal states (compare rows 1 and 6) and compensation for possible mismatch between the internal model and actual motion.

**Figure 6.**
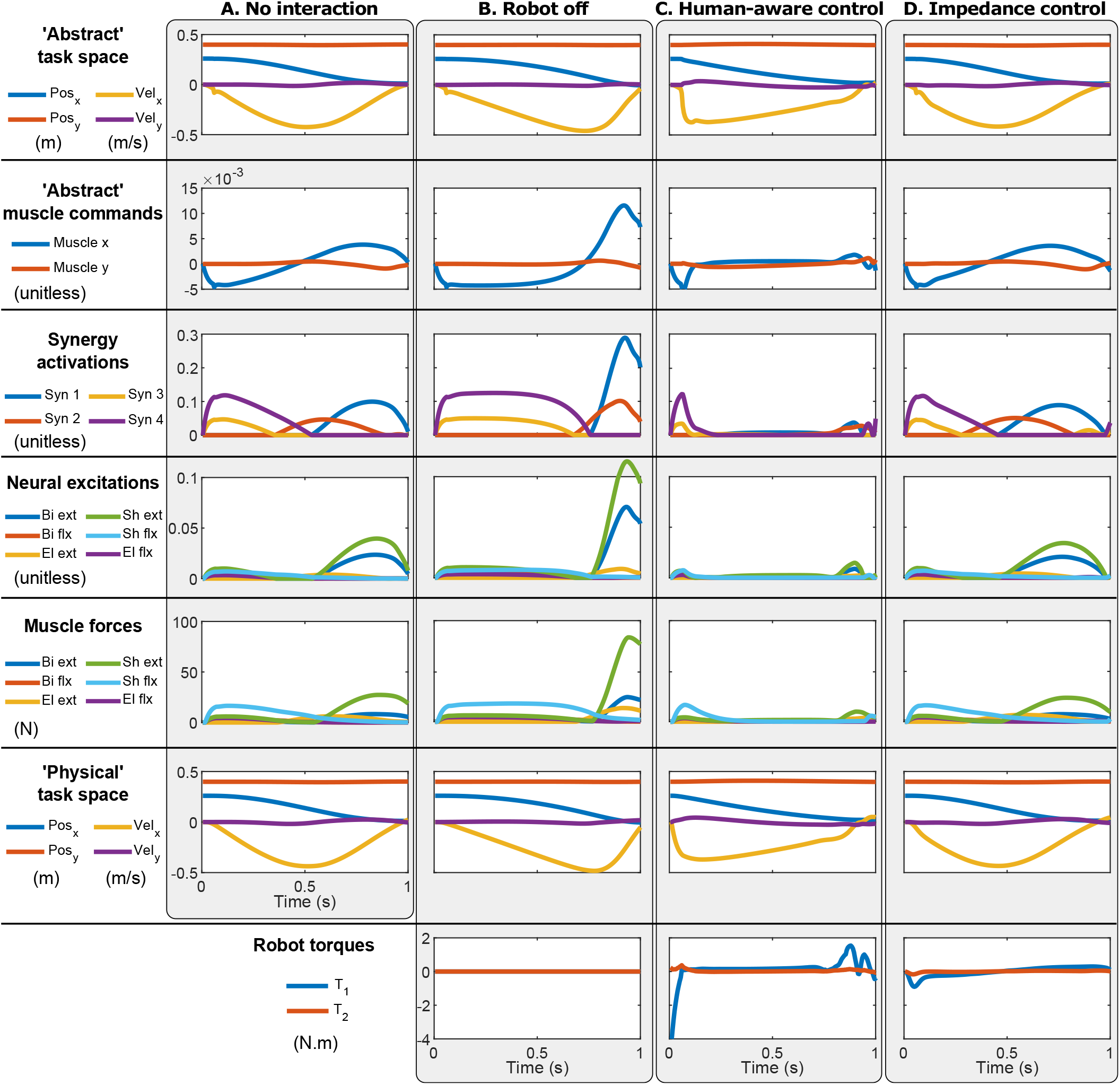
The simulation results. **A**. Human movement in isolation. **B**. Human+robot motion when the robot is turned off (zero motor torques). **C**. Human-robot interaction with the human-aware controller. **D**. Human-robot interaction with an impedance controller.

Interactions with an unactuated robot in the second simulation (Figure 6**B**) introduced a visible lag in movement (velocity peaked later) and increased muscular effort to accelerate and decelerate the added inertia of the robot. Despite the unmodeled dynamics of the robot, the arm arrived at the target because of its feedback structure.

The introduction of the human-aware controller in the third simulation significantly altered movement patterns (Figure 6**C**). In this movement, the human’s neuromuscular controller always wanted to compensate for the unknown interactions with the robot. On the other side of the interaction, the robot’s human-aware controller could predict these compensations and optimally adjust the interaction forces to minimize muscular effort, albeit at the cost of natural movement dynamics. Note the significant decrease in the neuromuscular activities (Figure 6**C**). This contrasted with the behavior that emerged from the impedance control in the fourth simulation (Figure 6**D**), where the robot merely followed the predetermined path without effectively collaborating with the user; the robot only carried its own inertia.

## 5 Discussion

In this paper, I presented a predictive model for human movement with two objectives in mind: the ability to predict movements with biologically realistic neuromuscular attributes and the ability to make such predictions faster than real-time for robot control purposes. I also presented a novel *human-aware control* paradigm, which employed the developed model for optimal control of human-robot interactions.

The holistic human movement model was truly predictive and only required the task description—in this case, target position and movement time—to predict the kinematics and neuromuscular activities similar to experimental data [37]. The model could also handle the unmodeled dynamics and disturbances and finish the tasks successfully. This holistic model could replicate human behavior faster than real-time; it took 350 *ms* to simulate 1.2 *s* movement time (on a laptop with Core i7 7660U CPU and 16 GB RAM, running MATLAB 2022b), which was significantly faster than other predictive models, e.g. [30]. Still, improvements to the model and controller’s computational efficiency are needed to make the human-aware controller solvable in real-time.

Although the holistic model remains to be rigorously validated against experimental data, individual modules within the model were based on well-supported biomechanical models and neuroscientific theories, contributing to the model’s validity as a whole. Specifically, the notions of internal models [13], optimal estate estimation [9], and optimal motor planning [9] are widely discussed in the computational neuroscience studied and are backed by experimental data. Further, the coordinated activity of muscles has been extensively documented [38], leading to realistic reconstruction of muscle activities using a small set of synergies. Lastly, the inclusion of the physiological characteristics of the musculoskeletal system [31] contributes to the bio-fidelity of the entire model. This holistic model, however, still misses important physiological attributes that shape human movement, e.g., reflex structures [39] and mechanical stiffness [40], which can improve predictions of in physical human-robot interaction, especially in response to sudden disruptions [18].

It must be noted that the user’s neuromuscular model was assumed to be *naive* to the robotic interactions, i.e., the user was not expecting any interaction force. However, the proposed human-aware control framework may include human-robot co-adaptation. The interaction forces from the robot can be modeled as a state-dependent force *F*_*robot*_ = **A**_*k*_**X**_*k*_ that may be *learned* by the human; its parameters can be included in the internal model (3) and iteratively updated using movement error [41]. Likewise, the robot’s control-oriented model can be updated iteratively as the user continues to react to the robot, e.g., using homotopy optimization [42].

The objective of the human-aware controller in this report was to minimize the user’s effort, which is only one of the many possibilities. Once the robot’s controller has access to a mathematical representation of individual aspects of the neuromuscular control system, e.g., the internal model’s states or the muscle synergies, the human-aware controller may seek to minimize any function of those variables; for instance, to increase or suppress the activation of a muscle synergy for motor learning and rehabilitation.

## 6 CONCLUSIONS

This paper presented a predictive model for human movement and a novel human-aware control paradigm for human-robot collaboration. Rooted in biomechanical and neuroscientific principles, the holistic model emulated human motor behaviors and enabled real-time prediction of neuromuscular activities. Leveraging this model, the human-aware control algorithm optimized robot actions to reduce user effort in interactive tasks, as validated in simulations with a robotic arm. These preliminary results reveal a promising approach to enhance human-robot collaboration, setting the stage for future research to refine and expand its real-world applications.

## References

[1] F. C. Anderson and M. G. Pandy, “Dynamic Optimization of Human Walking,” Journal of Biomechanical Engineering, vol. 123, no. 5, p. 381, 2001.

[2] A. J. Meyer, C. Patten, and B. J. Fregly, “Lower extremity EMG-to-moment modeling for walking with automated calibration of musculoskeletal geometry,” PLoS One, vol. 12, no. 7, pp. 1–24, 2017.

[3] A. L. Kinney, T. F. Besier, D. D. D’Lima, and B. J. Fregly, “Update on Grand Challenge Competition to Predict in Vivo Knee Loads,” Journal of Biomechanical Engineering, vol. 135, no. 2, p. 021012, Feb. 2013.

[4] T. K. Uchida, A. Seth, S. Pouya, C. L. Dembia, J. L. Hicks, and S. L. Delp, “Simulating Ideal Assistive Devices to Reduce the Metabolic Cost of Running,” PLOS ONE, vol. 11, no. 9, p. e0163417, Sept. 2016.

[5] M. Ezati, B. Ghannadi, and J. McPhee, “A review of simulation methods for human movement dynamics with emphasis on gait,” Multibody System Dynamics, vol. 47, no. 3, pp. 265–292, Nov. 2019, publisher: Springer Nature B.V. ISBN: 1104401909685.

[6] M. Febrer-Nafríia, A. Nasr, M. Ezati, P. Brown, J. M. Font-Llagunes, and J. McPhee, “Predictive multibody dynamic simulation of human neuromusculoskeletal systems: a review,” Multibody System Dynamics, vol. 58, no. 3-4, pp. 299–339, Aug. 2023.

[7] M. Ackermann and A. J. van den Bogert, “Optimality principles for model-based prediction of human gait,” Journal of Biomechanics, vol. 43, no. 6, pp. 1055–1060, 2010, publisher: Elsevier ISBN: 0021-9290.

[8] E. Todorov and M. I. Jordan, “Optimal feedback control as a theory of motor coordination,” Nature Neuroscience, vol. 5, no. 11, pp. 1226–1235, 2002, ISBN: 1097-6256 (Print)\r1097-6256 (Linking).

[9] S. H. Scott, “Optimal feedback control and the neural basis of volitional motor control,” Nature Reviews Neuroscience, vol. 5, no. 7, pp. 532–544, 2004.

[10] A. Q. Keemink, H. Van Der Kooij, and A. H. Stienen, “Admittance control for physical human–robot interaction,” The International Journal of Robotics Research, vol. 37, no. 11, pp. 1421–1444, Sept. 2018.

[11] S. F. Atashzar, M. Shahbazi, M. Tavakoli, and R. V. Patel, “A Passivity-Based Approach for Stable Patient–Robot Interaction in Haptics-Enabled Rehabilitation Systems: Modulated Time-Domain Passivity Control,” IEEE Transactions on Control Systems Technology, vol. 25, no. 3, pp. 991–1006, May 2017.

[12] B. Ghannadi and J. McPhee, “Optimal Impedance Control of an Upper Limb Stroke Rehabilitation Robot,” in ASME 2015 Dynamic Systems and Control Conference. Columbus, Ohio, USA: American Society of Mechanical Engineers, Oct. 2015, p. V001T09A002.

[13] D. McNamee and D. M. Wolpert, “Internal models in biological control,” Annual Review of Control, Robotics, and Autonomous Systems, vol. 2, no. 1, pp. 339–364, May 2019.

[14] E. Bizzi, F. A. Mussa-Ivaldi, and S. Giszter, “Computations underlying the execution of movement: a biological perspective,” Science, vol. 253, no. 5017, pp. 287–291, July 1991, ISBN: 0036-8075 (Print).

[15] J. Diedrichsen, R. Shadmehr, and R. B. Ivry, “The coordination of movement: optimal feedback control and beyond,” Trends in Cognitive Sciences, vol. 14, no. 1, pp. 31–39, Jan. 2010.

[16] S.-H. Yeo, D. W. Franklin, and D. M. Wolpert, “When optimal feedback control is not enough: feedforward strategies are required for optimal control with active sensing,” PLOS Computational Biology, vol. 12, no. 12, p. e1005190, Dec. 2016.

[17] F. Crevecoeur and S. H. Scott, “Beyond Muscles Stiffness: Importance of State-Estimation to Account for Very Fast Motor Corrections,” PLoS Computational Biology, vol. 10, no. 10, p. e1003869, Oct. 2014.

[18] R. Sharif Razavian, M. Sadeghi, S. Bazzi, R. Nayeem, and D. Sternad, “Body Mechanics, Optimality, and Sensory Feedback in the Human Control of Complex Objects,” Neural Computation, pp. 1–43, Mar. 2023.

[19] C. M. Harris and D. M. Wolpert, “Signal-dependent noise determines motor planning,” Nature, vol. 394, no. 6695, pp. 780–784, Aug. 1998.

[20] E. Todorov, W. Li, and X. Pan, “From task parameters to motor synergies: A hierarchical framework for approximately optimal control of redundant manipulators,” Journal of Robotic Systems, vol. 22, no. 11, pp. 691–710, 2005, ISBN: 0741-2223 (Print)\n0741-2223 (Linking).

[21] E. Todorov and M. I. Jordan, “A Minimal Intervention Principle for Coordinated Movement,” in Advances in Neural Information Processing Systems: Proceedings of the 2002 Conference, 2002, pp. 27–34, iSSN: 10495258.

[22] R. Sharif Razavian, N. Mehrabi, and J. McPhee, “A model-based approach to predict muscle synergies using optimization: application to feedback control,” Frontiers in Computational Neuroscience, vol. 9, no. October, pp. 1–13, Oct. 2015.

[23] R. Sharif Razavian, B. Ghannadi, and J. McPhee, “A synergy-based motor control framework for the fast feedback control of musculoskeletal systems,” Journal of Biomechanical Engineering, vol. 141, no. 3, p. 031009, Jan. 2019.

[24] R. Sharif Razavian, B. Ghannadi, and J. McPhee, “On the relationship between muscle synergies and redundant degrees of freedom in musculoskeletal systems,” Frontiers in Computational Neuroscience, vol. 13, Apr. 2019.

[25] R. Sharif Razavian, “A human motor control framework based on muscle synergies,” Ph.D. dissertation, University of Waterloo, 2017.

[26] D. M. Wolpert, J. Diedrichsen, and J. R. Flanagan, “Principles of sensorimotor learning.” Nature reviews. Neuroscience, vol. 12, no. 12, pp. 739–51, Dec. 2011, publisher: Nature Publishing Group.

[27] D. Liu and E. Todorov, “Evidence for the flexible sensorimotor strategies predicted by optimal feedback control,” Journal of Neuroscience, vol. 27, no. 35, pp. 9354–9368, 2007.

[28] B. Ghannadi, R. Sharif Razavian, and J. McPhee, “Configuration-dependent optimal impedance control of an upper extremity stroke rehabilitation manipulandum,” Frontiers in Robotics and AI, vol. 5, Nov. 2018.

[29] T. Van Wouwe, L. H. Ting, and F. De Groote, “An approximate stochastic optimal control framework to simulate nonlinear neuro-musculoskeletal models in the presence of noise,” PLOS Computational Biology, vol. 18, no. 6, p. e1009338, June 2022.

[30] N. Mehrabi, R. Sharif Razavian, B. Ghannadi, and J. McPhee, “Predictive simulation of reaching moving targets using nonlinear model predictive control,” Frontiers in Computational Neuroscience, vol. 10, no. 143, Jan. 2017.

[31] D. G. Thelen, “Adjustment of Muscle Mechanics Model Parameters to Simulate Dynamic Contractions in Older Adults,” Journal of Biomechanical Engineering, vol. 125, no. 1, pp. 70–77, Feb. 2003.

[32] B. Ghannadi, “Model-based Control of Upper Extremity Human-Robot Rehabilitation Systems,” Ph.D. dissertation, University of Waterloo, 2017, publication Title: PhD Thesis. University of Waterloo.

[33] F. Allgoöwer, R. Findeisen, and Z. K. Nagy, “Nonlinear model predictive control: From theory to application,” Journal of the Chinese Institute of Chemical Engineers, vol. 35, no. 3, pp. 299–315, 2004.

[34] F. Allgoöwer, T. A. Badgwell, J. S. Qin, J. B. Rawlings, and S. J. Wright, “Nonlinear Predictive Control and Moving Horizon Estimation — An Introductory Overview,” in Advances in Control, P. M. Frank, Ed. London: Springer London, 1999, pp. 391–449.

[35] C. Rao, J. Rawlings, and D. Mayne, “Constrained state estimation for nonlinear discrete-time systems: stability and moving horizon approximations,” IEEE Transactions on Automatic Control, vol. 48, no. 2, pp. 246–258, Feb. 2003, publisher: IEEE ISBN: 0018-9286.

[36] N. Hogan, “Impedance Control: An Approach to Manipulation: Part I—Theory,” Journal of Dynamic Systems, Measurement, and Control, vol. 107, no. 1, pp. 1–7, Mar. 1985.

[37] P. Morasso, “Spatial control of arm movements,” Experimental Brain Research, vol. 42, no. 2, pp. 223–227, Apr. 1981.

[38] E. Bizzi, V. C. K. Cheung, A. D’Avella, P. Saltiel, and M. C. Tresch, “Combining modules for movement.” Brain Research Reviews, vol. 57, no. 1, pp. 125–33, Jan. 2008.

[39] J. A. Pruszynski and S. H. Scott, “Optimal feedback control and the long-latency stretch response,” Experimental Brain Research, vol. 218, no. 3, pp. 341–359, 2012.

[40] E. Burdet, R. Osu, D. W. Franklin, T. E. Milner, and M. Kawato, “The central nervous system stabilizes unstable dynamics by learning optimal impedance,” Nature, vol. 414, no. 6862, pp. 446–449, 2001.

[41] J. Izawa, T. Rane, O. Donchin, and R. Shadmehr, “Motor adaptation as a process of reoptimization,” Journal of Neuroscience, vol. 28, no. 11, pp. 2883–2891, Mar. 2008.

[42] B. Ghannadi, R. Sharif Razavian, and J. McPhee, “A modified homotopy optimization for parameter identification in dynamic systems with backlash discontinuity,” Nonlinear Dynamics, vol. 95, no. 1, pp. 57–72, Jan. 2019.

